# Impact of gray matter signal regression in resting state and language task functional networks

**DOI:** 10.1101/094078

**Authors:** Luis C. T. Herrera, Gabriela Castellano, Ana C. Coan, M. Ludwing, C. L. César

## Abstract

A network analysis of the resting state (RS) and language task (LT) of fRMI data sets is presented. Specifically, the analysis compares the impact of the global signal regression of gray matter signal on the graph parameters and community structure derived of functional data. It was found that, without gray matter signal regression (GSR), the group comparison showed no significant changes of the global metrics between the two conditions studied. With gray matter signal regression, significant differences between the global (local) metrics for the conditions were obtained. The mean degree, the clustering coefficient of the network and the mean value of the local efficiency were metrics with significant changes. The community structure of group connectivity matrices was explored for both conditions (RS and LT) and for different preprocessing steps. When gray matter signal regression was performed, small changes of the community structure were observed. Approximately, the same regions were classified in the same communities before and after GSR. This means, that the community structure of the data is weakly affected by this preprocessing step. The modularity index presented significant changes between conditions (RS and LT) and between different preprocessing pipeline.

## 1. INTRODUCTION

In the present work, we studied how global signal regression of the gray matter signal changes brain network parameters and network community structure, when two conditions were compared. The two conditions under study were the resting state of the brain and a language (word production) task. Three main topics are relevant for the current work: 1) The effect of global signal regression (GSR) in fMRI data; 2) The resting state of the brain; 3) The functional differences between the brain at rest and the brain at task.

Regarding the first topic, a natural question that arises is, whether or not it is appropriate to apply GSR. This question has been focus of current scientific debate. Intrinsic to GSR, there exist unwanted and desirable consequences [1]. Among the unwanted effects of GSR we have that: i) It introduces negative correlations that were not present before [2–5]; ii) The correlation distribution is altered by the rescaling of the correlation values around a new mean. In consequence, it affects the values of local and long range correlations.; iii) It has negative effects for the interpretation of the group comparison [6]; iv) GSR reduces the spatial extent and intensity of coherent activated regions [7]. Among the desirable effects it is possible to enumerate: i) GSR removes uninteresting global fluctuations that blur the functional organization of the brain, and also increases the specificity of functional connectivity results. This is the case for the regression of the signal from ventricles and from white matter [8, 9]; ii) It increases the strength and reliability of experimental results. For example by taking whole head GSR, it is possible to improve the correspondence between functional and structural connectivity [8]; iii) GSR attenuates residual motion artifacts that can confound group comparison [10, 11].

The idea of using whole head GSR comes from the fact that the spatial distribution of the signal is present in a significant way in every gray matter voxel of the brain [8, 12]. Although this fact justifies the use of whole head GSR, as a suitable preprocessing step for fMRI-based brain network construction, the neuronal or non-neuronal origin of it, has not been well established. Many works using whole head GSR as a preprocessing step have been found that the brain posses a functional intrinsic architecture of anti-correlated networks [2, 13–15].

Regarding the second topic, the resting state of the brain, it has been established that the brain at rest presents significant correlations between BOLD-fMRI time series of anatomical regions separated from each other. These correlations have been interpreted as functional communication between brain regions, the so often called “resting state networks” [2, 14, 16]. The current view about resting state networks establishes that they constitute a baseline for brain activity. Brain (resting state or task) networks from fMRI data can be detected using model dependent algorithms such as the seed driven approach see, e.g.,[2, 12, 17], or using algorithms independent of models, such as Principal Component Analysis (PCA) [18] and Probabilistic Independent Component Analysis (PICA) [19]. More recently, the network science framework has been applied to the study of brain connectivity. The advantage of the network formalism is that offers a natural context to study in a quantitative way, the functional and anatomical features of the brain [20–25].

Finally, respect to previous research on functional differences between the resting state of the brain, and the brain in a guided task, the important conclusions obtained include: i) The functional architecture of the brain present in resting state is also present during many tasks [26, 27]; ii) The recognition of a primary and secondary network cores that shape the brain activity. One primary core that is stable, and common to many task states, and one secondary core that is flexible and transient [26–29]; iii) As consequence of the different configuration of the networks, the connectivity patterns can be used to identify the underlying task [28, 30]. The present work has some common and different characteristics with the previous work presented in [31]. Among the common characteristics, we have that the conditions studied in both works are the same, the resting state and a language task, also both works studied the functional connectivity patterns of the conditions. Among the characteristics that are different between the works, we have that, the present paper gave more attention to the study of the network metrics derived from the connectivity patterns, and the community structure of them.

Therefore, the aims of this work are: 1) Answer the question whether GSR should be applied as a preprocessing step in a graph analysis of brain networks from fMRI data; 2) Find out if network features can be used to distinguish between different brain conditions, namely, the resting state (RS) and a language task (LT).

The evidence presented reinforce the evidence that the GSR introduces global and local changes in the group comparison the metrics, and also suggest that the gray matter signal regression may be a suitable preprocessing step for the fMRI images.

**FIG. 1.**
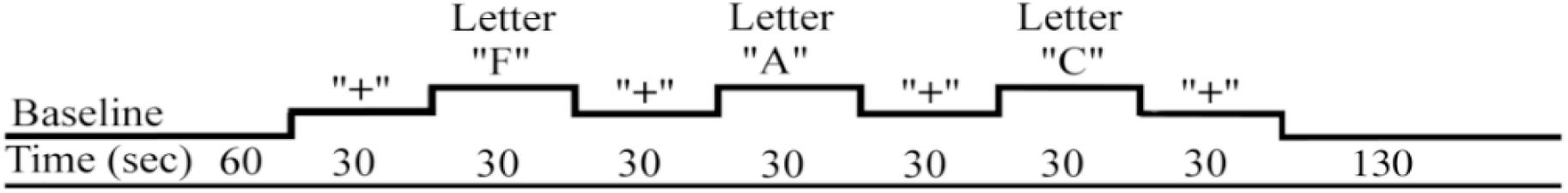
Language task paradigm. First 60 seconds (baseline period): black screen was shown to the subject. Cross (rest periods): black cross on white background. Letter (task periods): black letter on white background. Last 130 seconds (return to baseline): black screen.

## 2. MATERIAL AND METHODS

### 2.1. Subjects and protocols

Ten healthy subjects (mean age 35 ± 10, 6 women) participated in this study. The study was approved by the Ethics Committee of our institution, and all subjects signed an informed consent form prior to data acquisition. All subjects took part in two fMRI sessions (one RS and one LT) separated by a 6-minute anatomic scan. In the RS session, they were instructed to relax, to keep their eyes opened, to not fall asleep, and to think of nothing in particular, during 6.6 minutes. In contrast, in the LT session (also with a duration of 6.6 minutes) they were requested to focus their attention on a cross for 30 seconds (baseline condition), and next to silently think of as many words as possible beginning with the displayed letter, also for 30 seconds (experimental condition). The cross (‘+’) was presented in four blocks alternating with three different letters (‘F’, ‘A’, and ‘C’), presented in three different blocks Figure 1. This paradigm was the same as the one used in [32]. Stimuli were presented in a monitor positioned at the head of the magnet, and were visualized by the subject by means of a mirror attached to the head coil.

### 2.2. Acquisition Parameters

FMRI data were acquired with a 3 *T* Achieva magnetic resonance (MR) scanner (Philips, The Netherlands), using an EPI sequence with repetition time (*TR*) = 2000 *ms*, echo time (*TE*) = 30 *ms*, a voxel size 3 × 3 × 3.5 *mm*^3^ and 30 slices for the RS session, and an isotropic voxel size 3 × 3 × 3 *mm*^3^ and 40 slices for the LT session. An anatomic scan, consisting of a high resolution volumetric T1-weighted image, with voxel size 1 × 1 × 1 *mm*^3^ and *TR/TE* = 7000/3.24 *ms*, was also acquired, to be co-registered to the fMRI images.

**FIG. 2.**
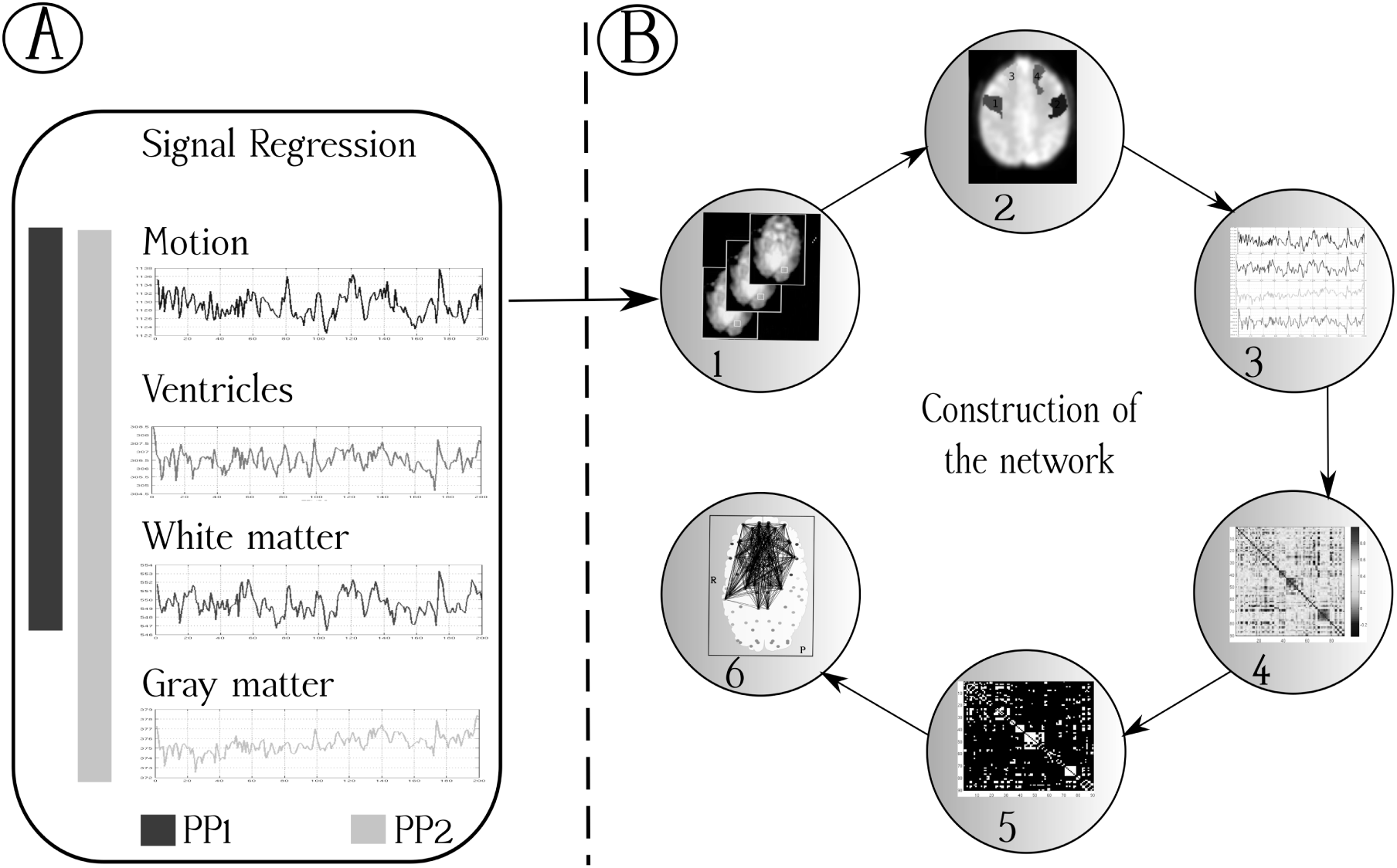
A) Different preprocessing pipelines (PPs) considered for network construction. PP1 consisted in signal regression of motion, ventricles and white matter. PP2 additionally regressed the mean gray matter signal. B) Scheme for the brain network construction. 1) Preprocessing of functional images; 2) Anatomical division of the brain; 3) Time series extraction; 4) Connectivity matrix construction; 5) Threshold definition for adjacency matrix construction; 6) Resulting undirected brain network.

### 2.3. Preprocessing

Image preprocessing was conducted using SPM8 (Wellcome Department of Cognitive Neurology, London, UK) running under MATLAB. The first five volumes of each participant were automatically discarded in the acquisition sequence. The preprocessing steps were: (1) correction of within-volume time differences between slices; (2) realignment of the volumes to the mean volume to correct for inter-volume movements; (3) co-registration of the anatomical image and the mean functional image; (4) spatial normalization of the functional volumes to a standard MNI template; (5) spatial smoothing with a Gaussian kernel of FWHM equal to double of the voxel size in each direction. A linear regression was used to remove the influence of head motion and of signals from the cerebrospinal fluid and white matter [9]. **In this paper, the term GSR will be used to describe the regression of the gray matter signal**. The impact of this preprocessing step on graph metrics and community structure was considered in many works [11, 33, 34]. To evaluate this effect, two type of preprocessing pipelines were considered, one with and other without GSR, identified as PP1 and PP2 respectively (see Figure 2). Finally, the fMRI data were temporally band-pass filtered in the range [0.01−0.08] Hz.

### 2.4. Network construction

For this section and the following, we consider a undirected non-weighted network represented as a graph *G(E, V)*, where *V* is a set of nodes, connected by a set of edges *E*. Figure 2 presents the methodology followed to construct the networks.

The set of nodes *V* are compose by the anatomical regions. Using the atlas developed by Tzourio Mazoyer et al., [35] the set of nodes are composed by 90 vertices corresponding to 90 anatomical regions (45 regions for each hemisphere). Subsequently, the mean time series of each region was extracted. Links were built by first calculating Persons correlation between all possible pairs of nodes *(i, j)*, considering the whole-time series. Note that although the LT paradigm contained task and rest periods, in this manner, we dealt with the temporal intervals in the same way for both RS and LT [36, 37]. Two nodes were classified as connected if the corresponding correlation coefficient was above a defined threshold *ρ*. The resulting network was then represented as a matrix A, known as the adjacency matrix, with elements given by a_*ij*_ = 1, if r_*i,j*_ > *ρ*, and a_*ij*_ = 0, if *r*_*ij*_ ≤ *ρ*, where *r*_*ij*_ was the Pearson correlation coefficient between the time series *x*_*i*_(*t*), *y*_*j*_(*t*) corresponding to nodes and *i* and *j*, given by

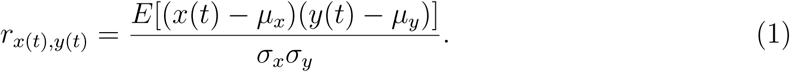

It is important to stress that we used only the positive values of the correlation coefficients to build the networks [21].

### 2.5. Network metrics

The undirected networks represented by the adjacency matrices *A* are functions of the correlation threshold *ρ*. A set of networks (i.e., adjacency matrices) were generated for different values of *ρ* within the interval [0.1 − 0.4], with increments of 0.1. In total, four networks were generated for each subject in each condition (RS and LT), with all networks fulfilling the requirement of being totally connected. From the networks four metrics were extracted, using the toolbox presented in [21]: 1) The degree of the node, which reflects the number of links each node has for a given threshold ( is the number of nodes, or regions):

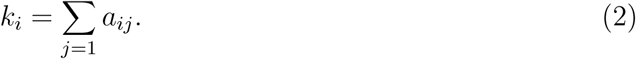

1. The cluster coefficient *c*_*i*_, which quantifies the density of connections around a node *i*:

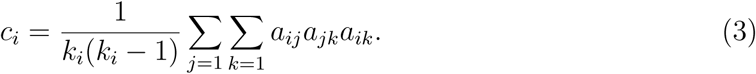
2. The local efficiency of the node *i*, which characterizes the efficiency of its neighbors in exchanging information once *i* is removed [38]

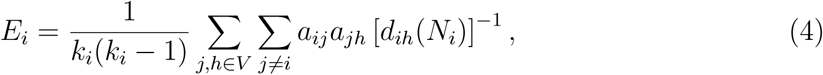

where [*d*_*ih*_(*N*_*i*_)]^−1^ is the shortest path between the nodes *j* and *h*, that contains only neighbors of the node *i*.
3. Characteristic path length *L*, that is constructed with the geodesic paths *d*_*ij*_ between the nodes *(i, j)* of the graph:

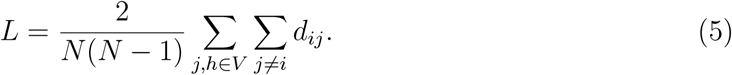

The first three metrics (degree, cluster coefficient and local efficiency) are local metrics, while the last (characteristic path length) is a global metric. From the local metrics, global metrics were computed, by averaging the values over all network nodes. The global metrics were studied with the intention of detecting global network changes for the different conditions.

Besides, analyzing local and global graph metrics, the correlation histograms were also computed, to verify possible differences in correlation values distributions resulting from different conditions (RS and LT) and pipelines.

### 2.6. Statistical analysis

To compare correlation histograms from different conditions and resulting from different preprocessing pipelines, the Kolmogorov-Smirnov test was used. To compare network metrics among these conditions and pipelines, a nonparametric statistical test (Wilcoxon test) was used, with a significance level of *p* ≤ 0.05. The Wilcoxon test was chosen due to the low number of samples (*N*=10). To control for the false discovery rate, the resulting p-values were corrected with the technique introduced by [39], and detailed in [40].

### 2.7. Group connectivity matrices and community detection

A group connectivity matrix *R*_*m*_*(i, j)* representative for each condition (*m* = *RS, LT*) was generated, using the individual connectivity matrices of the subjects. For this, the correlation coefficients for each subject, given by equation 1, were transformed to z-values using:

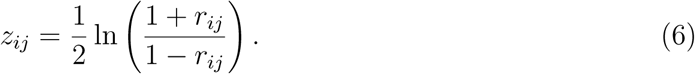

In this space (z), one average matrix *Z*_*m*_*(i, j)* was computed with the z-transformed connectivity matrices of the ten subjects [41]. Finally, the z-values in this matrix were transformed back into correlation coefficient values, using the inverse transform of equation 6, thus forming one connectivity matrix for each condition. The purpose of this construction was to make common patterns emerge from the data, because similar behaviors of the matrices will reinforce each other.

Before studying the community structure of the brain, we asked how different was the configuration of the temporal correlation in the two states. For this a scatter plot was made for the z-scored Pearson of the group matrices. In the x-axis was encoded the z-score of the temporal correlation between temporal series of two regions in the RS condition. In the y-axis was encoded the same information but for the LT condition. In this way, if a pair of regions presented high positive temporal correlation in RS and low positive temporal correlation in the LT, a point with high values (near to one) in the x-axis and low value (near to zero) in the y-axis would represent this configuration. The intention of this plot was to present an evidence that the temporal correlation values between the two states did not suffer extreme changes, indicating the existence of a common functional architecture [26, 27].

Relevant information was obtained from the community structure of the group matrices with the use of the modularity index [42]. The modularity index is a quantity defined under the assumption that networks can be divided in groups or communities. Then, if a given network can be divided in c communities, a symmetric matrix *e*_*ij*_ that represents the fraction of edges in the network that link vertices in the community *i* to vertices in the community *j* can be constructed. The dimension of the matrix is c × c. For an optimal community partition of the network, the trace of the matrix 
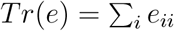
should possess a high value. The trace alone is not a good indicator of the quality of the partition. So another quantity *a*_*i*_ is defined as the row sum of the matrix 
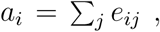
 and it represents the fraction of edges with at least one end in community *i*. In a network without a community structure we would have *e*_*ij*_ = *a*_*i*_*a*_*j*_. The modularity index is defined as next the difference

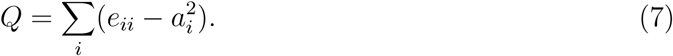

Values near to *Q* = 0 reflect networks with weak community structure, and values near to *Q* = 1 reflect networks with a strong community structure. Typical values of the modularity index for real networks, particularly brain networks, fall in the range of 0.3 to 0.7 [42, 43].

To explore the community structure information of the matrices, we used the adjacency matrices, for a correlation threshold *ρ* = 0. Many algorithms have been developed to detect the comunities [44, 45]. The partition of the nodes in communities was developed by the algorithm presented in [46], which maximizes the modularity index.

Finally, to visualize the performance of the community detection algorithm, the connectivity matrices were rearranged using the partition generated, and they were plotted in color scale like in [29, 43]. A distribution of the linear correlations, for the sets of nodes that belong to each community, was plotted in a boxplot for the LT and RS conditions and for each preprocessing pipeline.

## 3. RESULTS

### 3.1. Comparison between correlation histograms

According to the methodology described, representative BOLD time series for each anatomical region were computed using the mean of the voxel series inside each region.

**FIG. 3.**
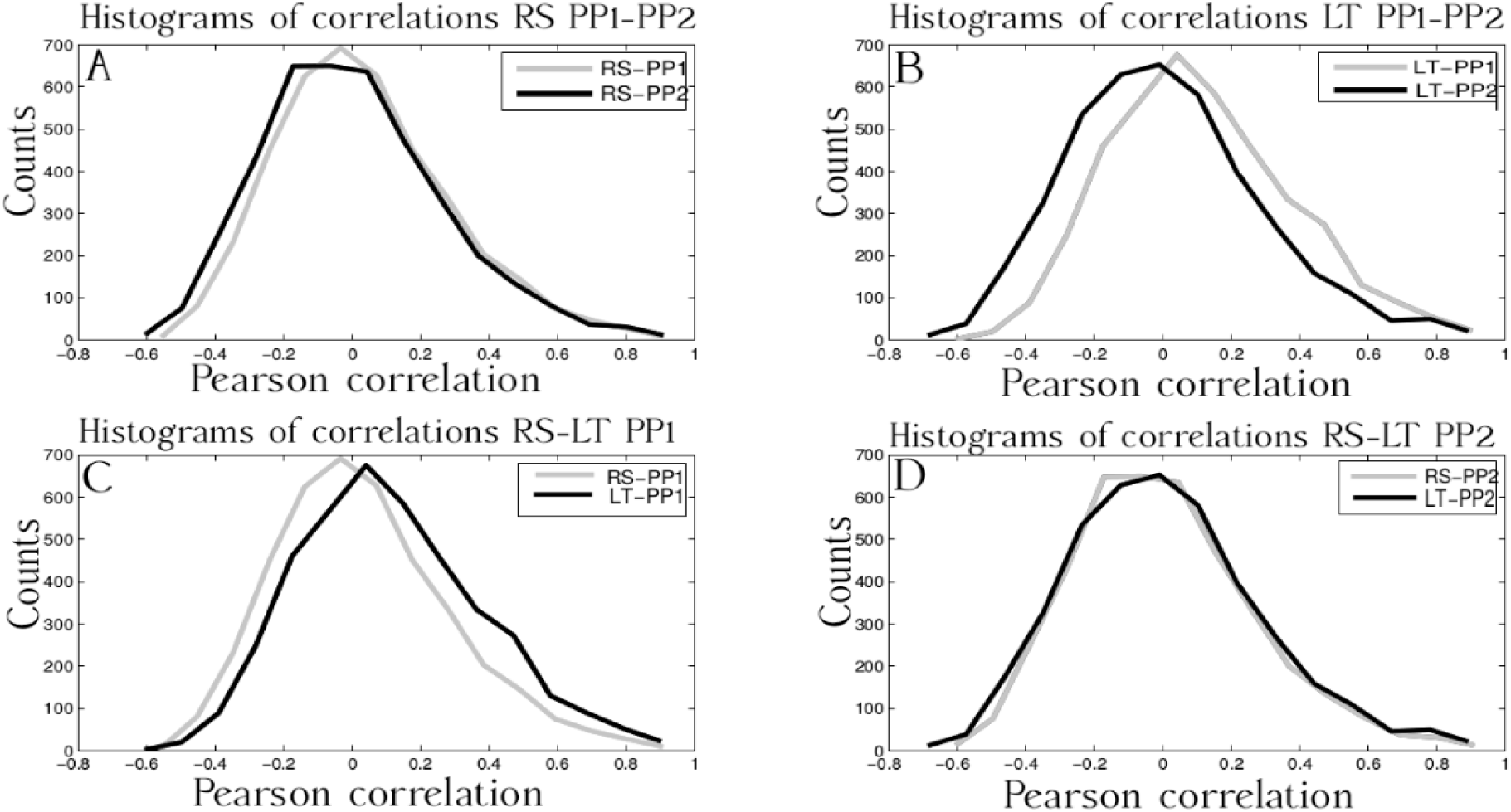
Histograms of correlations for the RS and LT conditions, for the different preprocessing steps studied, and for a representative volunteer. The figure A) present the histograms of correlations of the two preprocessing pipelines for the RS condition; B) Histograms of correlations for the LT, for the two preprocessing pipelines considered; C) Histograms of correlations for the LT and RS condition, for the PP1; D) Histograms for the RS and LT condition for the PP2. The Kolmogorov-Smirnov test shows no evidence, of statistical difference between them *p* > 0.05.

Pearsons correlation was computed for all pairs of time series. Figure 3 shows correlation histograms, for the two conditions and the two preprocessing pipelines, for one representative subject. Histograms generated using the same pipeline but for different conditions (Figures 3C and 3D) showed no statistical difference for any subject (*p* > 0.05, Kolmogorov-Smirnov test). Similarly, histograms generated with different pipelines for the same condition (Figures 3A and 3B) showed, as expected, an increase in negative correlations [2–4] for the GSR pipeline (PP2), these differences were not significant for any subject (*p* > 0.05, Kolmogorov-Smirnov test) with the use of the AAL.

**FIG. 4.**
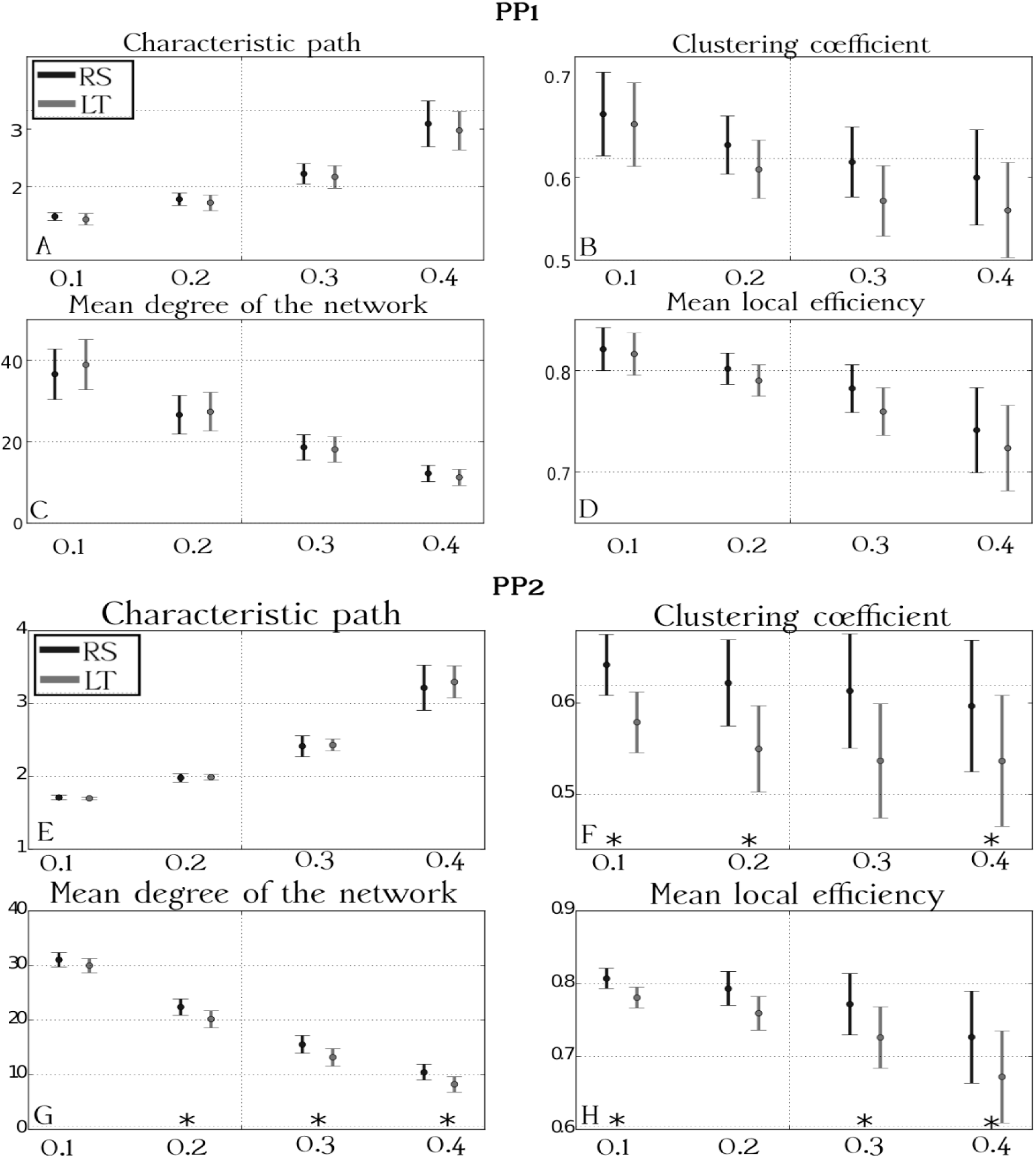
Average global metrics obtained over the 10 subjects for PP1 (top) and PP2 (bottom) in the two conditions: RS (black) and LT (gray). Error bars represent standard deviation over subjects. The horizontal axis shows the different positive thresholds used to construct the adjacency matrix: *T*1 = 0.1, *T*2 = 0.2, *T*3 = 0.3, *T*4 = 0.4. The figures A) and E) present the characteristic path length; The figures B) and F) present the clustering coefficient; The figures C) and G) present the mean network degree; The figures D) and H) present the mean value of the local efficiency of the network. Thresholds indicated with ‘*’ presented FDR corrected *p* – *values* < 0.05.

### 3.2. Global metrics

Figure 4 (top) shows plots for the global metrics averaged over the 10 subjects in the two conditions, using PP1 (without GSR, top part of the figure) and PP2 (with GSR, bottom part of the figure). Error bars indicate standard deviations over subjects. Four different thresholds indicated as T1, T2, T3 and T4 represent *ρ* = 0.1, *ρ* = 0.2, *ρ* = 0.3 and *ρ* = 0.4 respectively. Thresholds marked with a ‘*’ indicate statistical significance. The RS condition is represented in black and LT in gray. From the top of this figure, we see that there were no significant differences between conditions for any global metric (or threshold) with this preprocessing pipeline (*p* > 0.05, Wilcoxon test, FDR corrected). Differently than for the pipeline without GSR, we see from the bottom of Figure 4 that by applying GSR three of the global metrics (clustering coefficient, mean network degree and global efficiency) were significantly different among conditions, for at least three of the applied thresholds (*p* < 0.05, Wilcoxon test, FDR corrected). These metrics were all higher for the RS compared to the LT condition.

### 3.3. Local metrics

Given that PP2 showed significant differences between conditions for the global metrics, the corresponding local metrics were analyzed with the objective of identifying the set of regions that caused the observed global changes with this preprocessing pipeline. The metrics analyzed were thus the degree, the clustering coefficient and the local efficiency of the nodes. We chose to look only at threshold ρ = 0.4, given that this threshold presented significant changes between the conditions for all the metrics mentioned. For every metric, p-values were obtained using the Wilcoxon test, for each region. Subsequently, the FDR controlling statistics was applied, for controlling rates of *q* = 0.05, *q* = 0.1, and *q* = 0.2, as described and suggested in [40]. For the first two controlling rates, no significant changes for the local metrics between the different conditions were found. For the controlling rate *q* = 0.2 (Figure S1), we obtained sets of regions that presented significant changes of the metrics between the conditions.

The Table I presents the list of regions that showed significant changes (*p* < 0.2, Wilcoxon test, FDR corrected) for the local metrics: node degree, node clustering coefficient and local efficiency. All regions in this table, with exception of the Frontal-Mid-L region for the degree, presented higher values of the metrics (averaged over subjects) for the RS compared to the LT condition. This result supports the findings of the global metrics presented in Figure 4, where the global values of the metrics were higher in the RS condition than in the LT. With the intention to test if the results presented by the local metrics agreed with functional areas relevant for the task, the Online Brain Atlas Reconciliation [47] was used to identify areas in the AAL atlas that map the Broca (Brodmann’s areas 44L and 45L) and Wernicke’s areas (Brodmann 22L). Table II presents that list of regions; it also contains the local metrics that presented differences between the two states studied.

**TABLE I.** List of regions with significant changes (*p* < 0.2, Wilcoxon test, FDR corrected) of local metrics between the RS and LT conditions. Bold font indicates regions that presented significant changes in all three local metrics.

| Degree | Clustering coefficient | Local efficiency |
| --- | --- | --- |
| Frontal-Mid-L | **Frontal-Mid-Orb-R** | **Frontal-Mid-Orb-R** |
| Rolandic-Oper-R | Rolandic-Oper-L | Frontal-Inf-Orb-L |
| Olfactory-L | Insula-L | Rolandic-Oper-L |
| Olfactory-L | ParaHippocampal-L | Hippocampus-L |
| Frontal-Sup-Medial-R | Cuneus-R | Occipital-Mid-L |
| Frontal-Mid-Orb-L | Occipital-Sup-R | Occipital-Inf-L |
| **Frontal-Mid-Orb-R** | Occipital-Mid-L | **Fusiform-L** |
| Rectus-L | Occipital-Mid-R |  |
| Rectus-R | Occipital-Inf-L |  |
| ParaHippocampal-L | **Fusiform-L** |  |
| ParaHippocampal-R |  |  |
| Amygdala-R |  |  |
| Lingual-L |  |  |
| Occipital-Sup-L |  |  |
| Occipital-Sup-R |  |  |
| **Fusiform-L** |  |  |
| Fusiform-R |  |  |
| Caudate-R |  |  |
| Heschl-L |  |  |
| Temporal-Sup-L |  |  |
| Temporal-Pole-Mid-L |  |  |
| Temporal-Inf-R |  |  |

**TABLE II.** List of regions in the AAL atlas that compose the Broca and Wernieck’s areas. The list of regions were taken from the Online Brain Atlas Reconciliation [47]. Regions in the AAL that presented differences for the local metrics, between the two states are indicated by: 1) DEG, for degree of the node; 2) CLU, for cluster coefficient; 3) LE, for local efficiency.

| BA 44L- BA45L (Broca's area) - Local metric | BA 22L (Wernicke's area)-Local metric |
| --- | --- |
| Frontal-Inf-Oper-L - (LE) | Temporal-Mid-L |
| Front-Inf-Trian-L | Temporal-Sup-L - (DEG) |
| Insula-L - (CLU) | Temporal-Pole-Sup-L |
| Rolandic-Oper-L - (LE) | Heschl-L - (DEG) |
| Precentral-L | Angular-L |
| Frontal-Inf-Orb-L - (LE) | Rolandic-Oper-L - (LE) |
|  | Supramarginal-L |

**FIG. 5.**
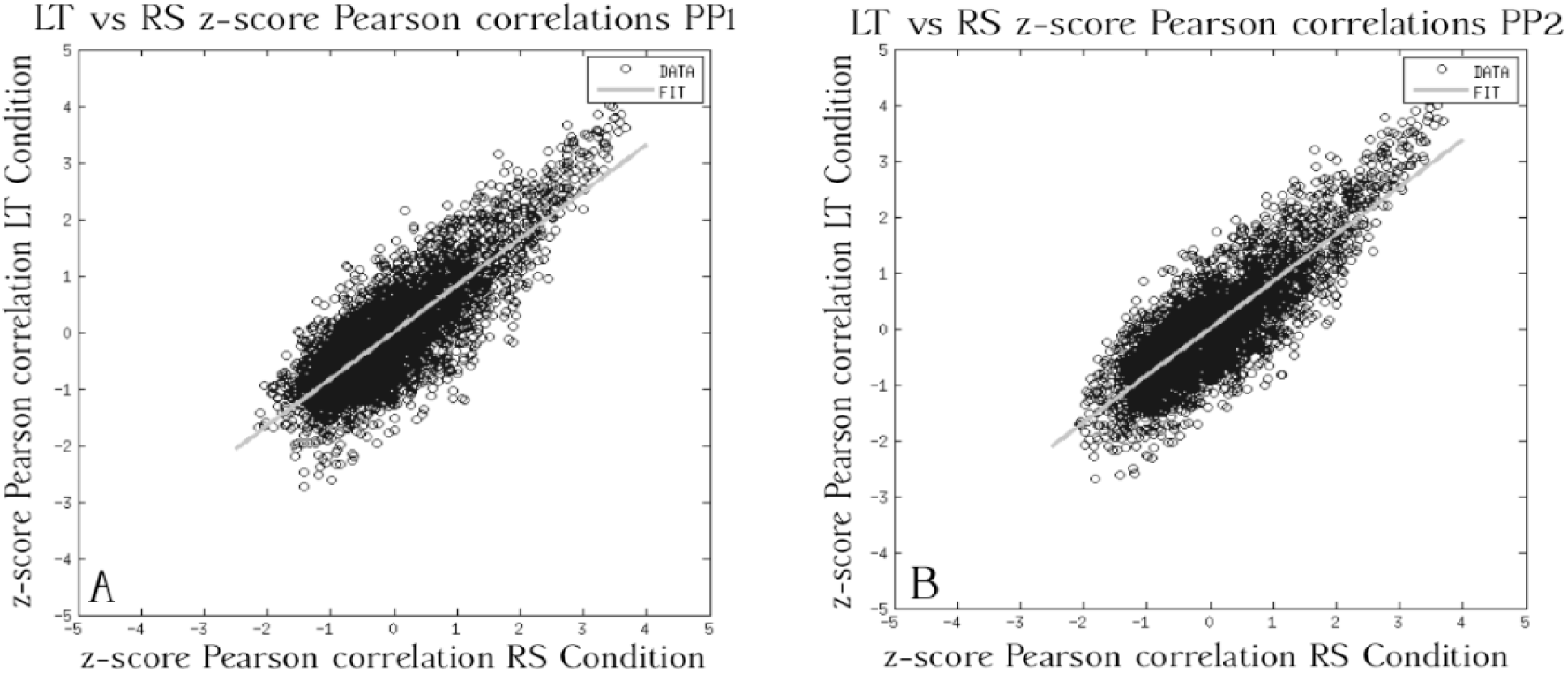
Scatter plot of the z-scored Pearson correlations in the two conditions presents linear dependency between them. A) Scatter plot for the z-scored Pearson correlations for the PP1; B) Scatter plot for the z-scored Pearson correlations for the PP2

### 3.4. Group connectivity matrices and community detection

In Figures 5A and 5B, as explained in Section 2.7, we present the scatter plot of the z-scored temporal correlations in RS and in LT. As expected, temporal correlations in the two states remained with similar values, that is low (positive or negative) temporal correlations in RS were obtained also in LT, and high temporal correlations (positive or negative) in RS were also obtained in LT. This fact supports the claim that the brain works with a basic functional architecture present in the two states [26, 27]. The model was fitted using the least squares method, showing statistical evidence (*p* < 0.05, t-test) that supports the existence of linear dependency between pairwise correlations in the two conditions [26].

**FIG. 6.**
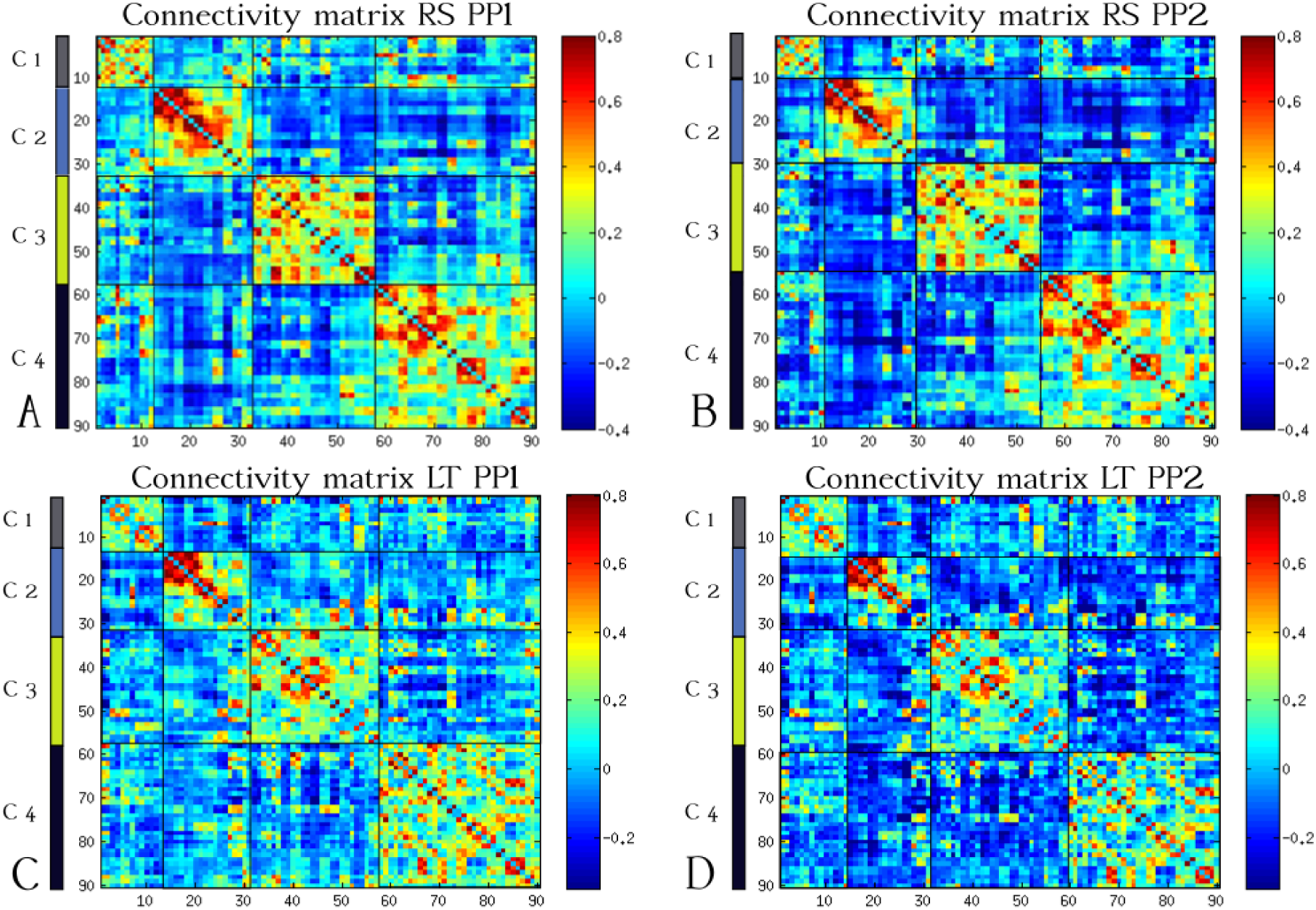
Group connectivity matrices for: A) RS condition with PP1; B) RS condition with PP2; C) LT condition with PP1; D) LT condition with PP2. The matrices were rearranged using the detected communities, ordered from the smallest to the largest.

### 3.5. Group connectivity matrices and community detection

The community detection for each of the group matrices was performed with the methodology presented in Section 2.7, and each connectivity matrix was reordered according to the communities found. Four communities were found for each matrix. Figure 6 shows the reordered group matrices for the RS and LT conditions, for each preprocessing pipeline, PP1 and PP2. The tables S1 to S8, in the Supplemental Materials present the communities found for each group matrix for the RS and LT condition respectively. To give an idea of the communities structure, Figures 7 and 8 show the communities in the RS and LT condition, respectively, for PP2. In these figures, the left side shows an axial representation of the brain, while the right side shows a sagittal view.

Figure 9 presents a plot of the mean value of the modularity index across subjects in each condition and preprocessing pipeline, with error bars representing the standard deviation over subjects. The preprocessing steps of the images alters significantly the modularity value. Higher modularity indexes were found for PP2 compared to PP1, and the RS condition presented higher modularity values compared to the LT condition.

Network modules are defined as a set of brain regions that are strongly connected to each other and weakly connected to the rest of the brain [29]. Then, with the intention of providing a statistical assessment for each community, the correlation distribution for each community in each condition and preprocessing pipeline is presented in Supplementary Figure S2. We obtained correlation distributions significantly different from a Gaussian distribution with zero mean for each community.

## 4. Discussion and conclusions

The work presented by us reinforce the evidence that the GSR introduces global and local changes in the group comparison the metrics, and also suggest that the gray matter signal regression may be a suitable preprocessing step for the fMRI images. The main goal of this work was to assess how GSR affects brain networks derived from fMRI data; particularly, how it affects the comparison between networks corresponding to different brain states, namely, the resting state and a task state, in this case consisting of a language task. Networks for these two states (or conditions) were therefore built following two preprocessing pipelines, one without (PP1) and another with (PP2) application of GSR. From these networks, we analyzed the correlation histograms, global and local network metrics, and communities.

The correlation histograms showed a visible, however non-significant, increase in negative correlations (shift to the left) when GSR was applied, which is in accordance to [8].

It was found that without applying GSR, it was not possible to distinguish between the RS and LT conditions using global network metrics. Interestingly, when introducing GSR, significant differences among global network metrics (mean degree, mean clustering coefficient and mean local efficiency) between RS and LT conditions were found. RS presented higher values of these metrics compared to LT, as expected. The mean degree represents the average number of connections of a region, i.e., to how many other regions it is functionally synchronized. Therefore, this result means that more regions are connected (or work together) in the resting state, when the brain is not focused in any specific task, than when the brain is actually performing a very specific task such as language, that requires that a smaller set of specific regions work together. The network clustering coefficient measures something somehow similar to the mean degree, which is how connected (in average) are the neighbors of a node. We see again that the decrease of the clustering coefficient for a specific task compared to the resting state makes sense, given that fewer regions are used in the former case. Finally, the mean local efficiency gives a measure of how well, information can travel across the network. Again, since for the language task only a few regions are recruited, other regions are “shut down” and the information does not reach them anymore, which would explain the decrease in efficiency.

**FIG. 7.**
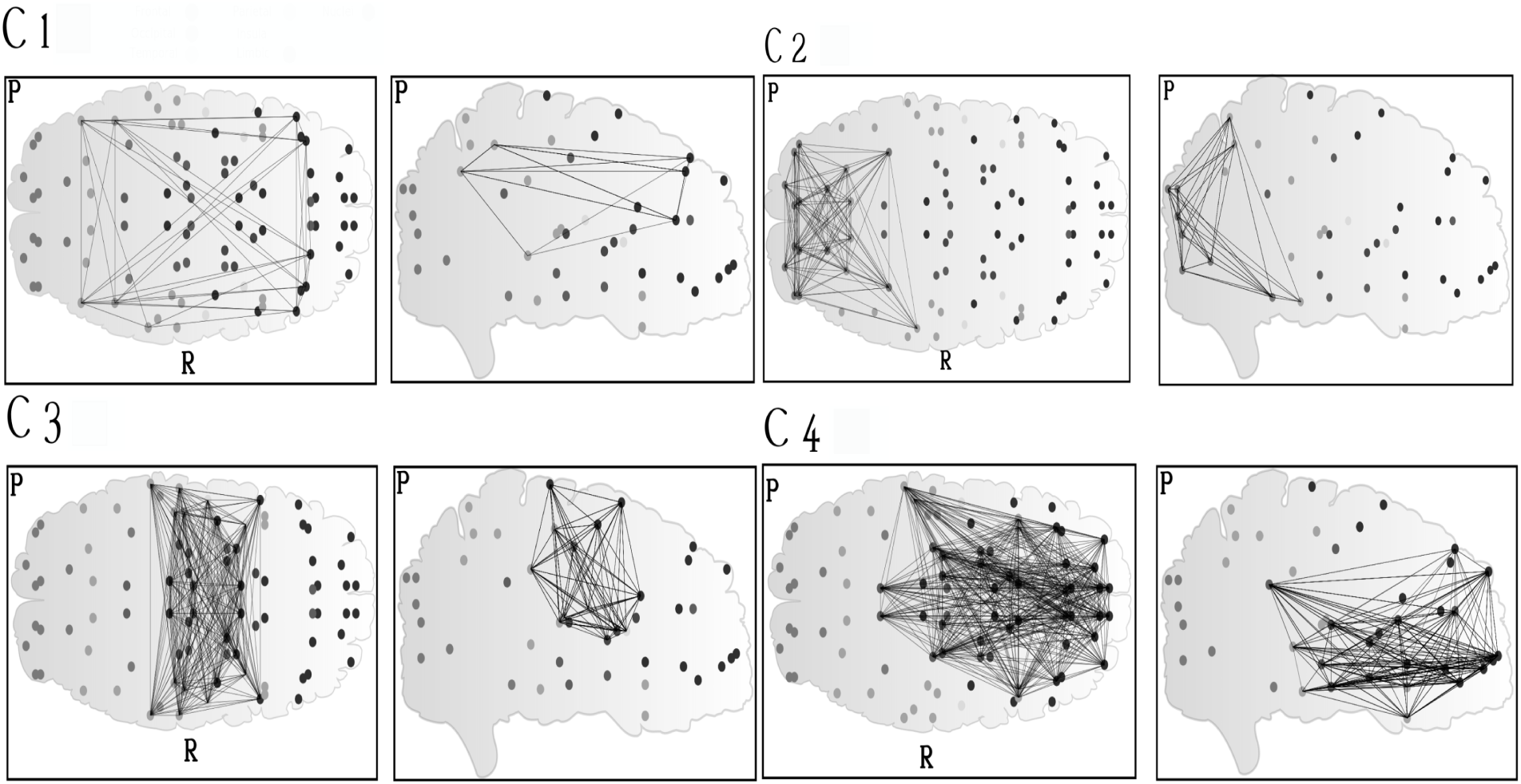
Communities detected for the RS condition with PP2 (see Tables S1-S4 right). Left: axial view, with right hemisphere indicated by the letter R and posterior regions with the letter P. Right: sagittal view.

**FIG. 8.**
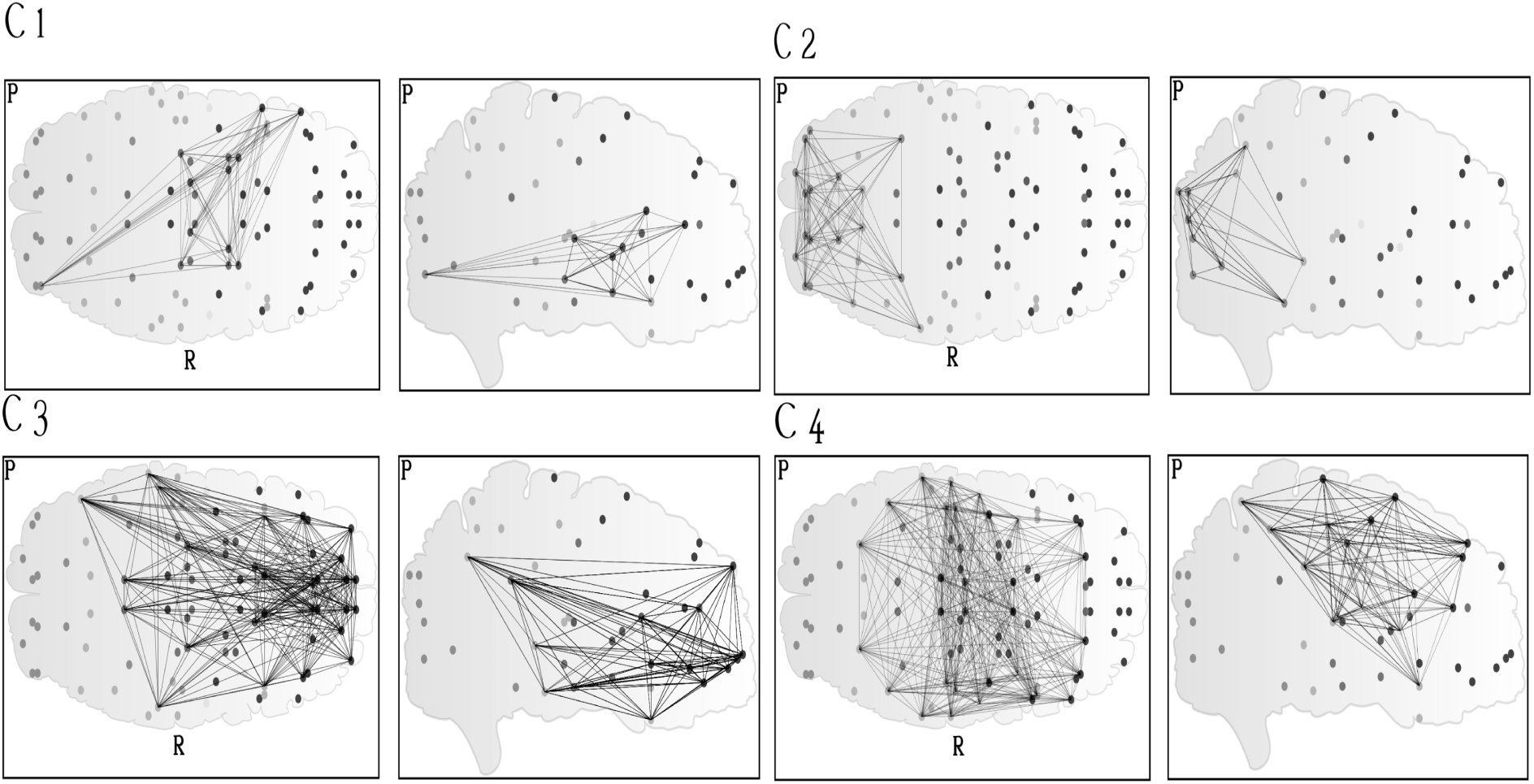
Communities detected for the LT condition with PP2 (see Tables S5-S8 right). Left: axial view, with right hemisphere indicated by the letter R and posterior regions with the letter P. Right: sagittal view.

**FIG. 9.**
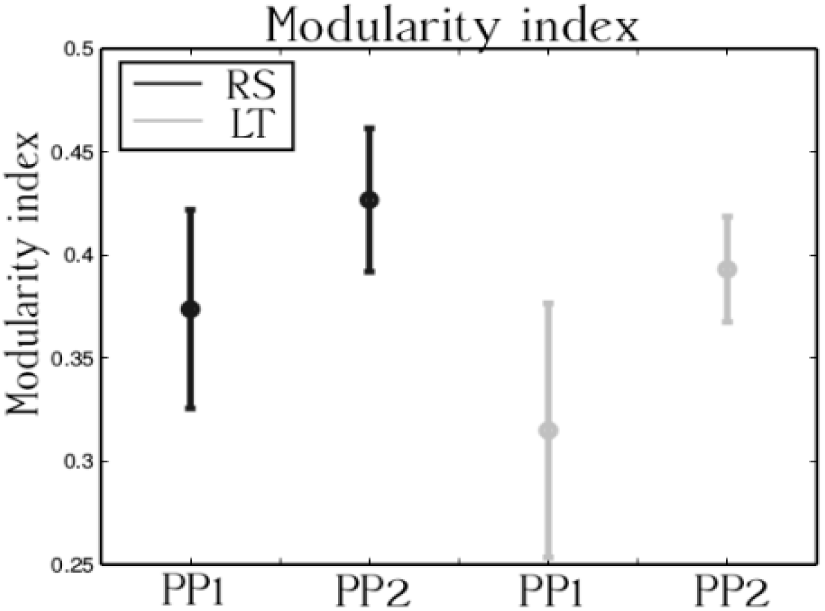
Modularity index equation (7) for the different conditions, and preprocessing pipelines, the error bars represent the mean and standard deviation for the 10 volunteers. The modularity index is modified by the different kind of preprocessing steps of the images. The PP2 produces an increase of this index when compare to the images that suffer the preprocessing PP1. For the same preprocessing pipeline, the RS condition shows higher values of the modularity than in the LT, this means that the community structure is strongest in RS.

In the work by Fox and colleagues [8], it was pointed out that the global signal of the whole head was present in a significant way in every voxel of gray matter. By the evidences provided in the present study, it seems that this signal obscures significant differences of the global network metrics between the two states. Nevertheless, it is possible that the obtained global differences could be spurious. In that case the study of the local metrics can help to provide evidence of the nature of the differences. If the local metrics reveal differences in areas that agree with the current task, this may suggest a certain degree of reliability for the global differences obtained.

The analysis with local metrics for PP2 allowed identification of the regions responsible for the global changes seen previously. In spite of the fact that only a relatively loose significance level allowed identification of these regions (*p* < 0.2, Wilcoxon test, FDR corrected), the regions found (Table I) make sense when considering that a language task was involved. In particular, 22 regions were found that changed their degree value from one condition to the other, 10 regions that changed their clustering coefficient, and seven that changed their local efficiency (Table I). The first region of the degree list Left frontal middle gyrus, is the only region that has, in average, higher degree value in the LT condition than in the RS condition. This indicates that this region increases its temporal correlation with other regions in the language task compared to the resting state. The other regions presented, in average, higher degree in the RS condition. The frontal medial left area is, indeed, involved with a passive intention of the subjects to move the mouth and say the words that they think. There were only two regions that presented changes in all three local metrics, namely, Left frontal middle gyrus orbital portion and left fusiform gyrus. The left frontal middle gyrus orbital portion area is involved with the intention to move the mouth to say the words of the task. The left fusiform gyrus is involved with word recognition [48].

As mentioned in the introduction, the present work shares a similarity with the work presented in [31], because there, a language task was compared with the resting state of the brain. In the mentioned work, it was found an increase in the correlation between the Broca and Wernicke’s areas in the language task condition, when compared to the resting state condition. In the present work, it was found that 7 of 13 regions that compose the Broca and Wernicke’s areas in the atlas AAL presented local changes of some of the studied metrics (node degree, clustering coefficient or local efficiency) between the conditions (Table II).

The Broca’s area (BA 44L- BA45L) is composed in the AAL by the: 1) left inferior frontal gyrus, opercular portion; 2) left inferior frontal gyrus, triangular portion; 3) left insula; 4) left Rolandic operculum; 5) left precentral gyrus, 6) left inferior frontal gyrus, orbital portion. Wernicke’s area (BA 22L) is composed by: 1) left middle temporal gyrus; 2) superior temporal gyrus; 3) temporal pole; 4) left Heschl gyrus; 5) left angular gyrus; 6) left Rolandic operculum; 7) left supramarginal gyrus [47]. The findings suggest that GSR of the gray matter signal helps to reveal local differences of the metrics between the two states. The differences appear in regions that agree with the language task in course.

In summary, we found that the network metrics studied were affected by GSR. When GSR was applied, some global and local metrics presented significant changes between the two conditions studied. Despite the low sample size (10 subjects), it was possible to identify regions with significant metric changes between the conditions.

Considering the distribution of the temporal correlations, we found that the scatter plot of the correlations between the two conditions presented evidence of a linear dependency between them, for the two preprocessing pipelines considered (Figure 5). In other words, in most cases, if two brain areas possess a low temporal correlation in the RS condition, they also possess a low temporal correlations in the LT condition. The same fact was confirmed for high positive and high negative temporal correlations. This agrees with the claim that there exist small changes in the functional architecture of the brain between different states [26, 27].

The spatial distribution in the brain of the connectivity architecture can be studied looking at the community structure of the data. For the same condition but with different preprocessing steps, each condition presented small changes in its community structure (see Tables S1 to S8). In the RS, four regions changed their community labels after GSR. For the LT condition, three regions changed their community labels after GSR.

The results may not be accurate because the use of the AAL atlas blurs the boundaries between functional areas and leads to poor network estimation [49, 50]. The group connectivity matrices were constructed with the objective of reinforcing coherent activities across the subjects and in this way overcome the potential signal blur between regions.

The first community of the RS condition may reflect the default mode network because this community includes the angular gyrus (left and right) and medial frontal cortex (left and right). The first community in the LT seems to reflect the underlying language task, because it involves connections between visual areas and areas that belong to the Broca and Wernickes areas such as the opercular and triangular portions of the left inferior frontal gyrus, and the temporal pole (see Tables S1 and S1).

The second community is similar for both RS and LT and includes the visual network (V1 and V2) and its connections with the inferior temporal and parietal regions. The primary visual network is known for its high level of connectivity, what explains its prominence in both conditions [51].

Other prominent network, the sensoriomotor network, was detected in third community of RS condition. The third community of LT and fourth community of RS conditions share similarities, and they include areas involved in executive and memory process, such as bilateral dorsolateral prefrontal cortex, hippocampus, parahippocampal gyrus and bilateral temporal lobes [30]. Finally, the fourth community of the LT condition involves both primary language areas and regions associated with the motor process of language and it can be also related to the task performed [52].

Considering the modularity index in the same condition (RS or LT), but for different preprocessing pipelines, it was shown an increase on its value when GSR was applied. Considering the same preprocessing pipelines but in different conditions, in RS the modularity index was higher than in LT. This means that the community structure is stronger in the RS condition. The values of modularity found in this work fall in the range of typical modularity values for brain networks, reported in previous works [43].

It is important to mention that in this work, only positive correlations were considered to construct the brain networks. This could be problematic, considering that GSR introduces negative correlations. In this way, the obtained metrics may not reflect the whole network change after GSR. One characteristic that remained unchanged with and without GSR was that the structure of the connectivity matrices (Figure 6) revealed that the interaction between the different network modules was primarily via negative correlations.

The findings provided in the present work support the claim that the global signal regression of the gray matter signal is a suitable preprocessing step for fMRI images, since this preprocessing step does not alter the community structure of the data, and reveals differences in the global and local metrics that involve areas engaged in the underlying task.

## Acknowledgements

This work was supported by the Brazilian governmental funding agencies: FAPESP (São Paulo Research Foundation, grants 2013/07559-3 and 2013/00099-7), CNPq (National Council of Scientific and Technological Development) and CAPES (Coordination for the Improvement of Higher Education Personnel). Special acknowledgments to Guilherme Beltramini, Marcus Aguiar, Edwin Forero, Rickson Mesquita, Jose Zapata, Jaroslav Hlinka and Elvis Lira for the academic discussions that enriched the present work. This work was also developed with resources from Instituto Nacional de Fotônica Aplicada à Biologia Celular INFABIC (CNPq grant 57391/2008-0, FAPESP grant 08/57906-3) and Biologia das Doenças Neoplásticas da Médula Óssea (FAPESP grant 11/51959-0).

